# Observational satiety: watching yourself eat induces more fullness than watching another

**DOI:** 10.64898/2026.08.28.747712

**Authors:** Juri Fujiwara, Hitoshi Kubo, Satoshi Kida

**Author notes:** **Corresponding author:** Juri Fujiwara, Department of Systems Neuroscience, Fukushima Medical University, Fukushima 960-1295, Japan.

## Abstract

Appetite is shaped not only by physiological need but by the sensory experience of eating and its social context. Can watching food being eaten itself induce satiety, and does it matter who is seen eating? In a functional MRI study (N = 41), participants watched videos of a wanted snack being eaten from their own perspective (self) or, using identical footage shown vertically inverted, as another’s (other), holding food identity and visual content constant so that only the attributed agent varied. Observed eating reduced wanting, but not liking, for the eaten foods, whether one’s own or another’s; crucially, reported fullness increased only when the eating was seen as one’s own. In the brain, food-value regions responded to watching eating in both conditions, a shared signal that strengthened over time. Yet only self-attributed eating engaged a self-specific, value-related response in the ventral striatum and medial prefrontal cortex, and orbitofrontal activity during self-eating scaled with each person’s reported satiety, whereas watching another eat instead engaged the temporoparietal mentalizing network without a comparable rise in fullness. A large online survey (N = 1,000; ages 15–97) reproduced the behavioral effect across the adult lifespan, and a real-eating experiment reproduced its sensory-specific pattern. Watching eating, and whose eating it is, can thus recalibrate appetite through separate food-value and social-cognitive routes. This “observational satiety” offers a non-invasive route to study food wanting, of potential relevance to social eating and to today’s food-media environments.

**Highlights:**

- Watching yourself eat induced satiety, without any eating
- This satiety required the eating to be seen as one’s own, not another’s
- A self-specific value-related response appeared in ventral striatum and medial PFC
- Watching another eat instead engaged mentalizing regions (TPJ, dmPFC)
- A large survey (N = 1,000) and a real-eating study showed the same behavioral effect

## 1. Introduction

Appetite is not driven by physiological energy needs alone. It also depends on sensory experience, expectation, memory, and the social context of eating. A familiar example of this flexibility is sensory-specific satiety: after eating a particular food, its pleasantness declines while foods that have not been eaten remain appealing. This is the everyday reason there is still ‘room for dessert’ after a filling savory meal. Such selective satiety reflects the sensory experience of eating rather than metabolic feedback (B. J. Rolls et al., 1981; B. J. Rolls, 1986; Hetherington et al., 1989). It has a well-documented basis in the brain’s food-value system, which assigns value to food and turns it into the motivation to eat (Bartra et al., 2013; E. T. Rolls, 2015), and this system is engaged even by the mere sight of food, before any eating (van der Laan et al., 2011). To describe appetite change precisely, we use the standard distinction in appetite research between two components of food reward: the current motivation to eat a food (incentive value, or ‘wanting’) and the sensory pleasure it gives (‘liking’), which are dissociable and can change independently (Berridge, 2009; Finlayson et al., 2007). A selective decrease in wanting while liking is unchanged therefore reflects a change in a food’s motivational value rather than in how much it is liked. Throughout, ‘satiety’ refers to the subjective feeling of fullness that participants rate directly, not to the decrease in wanting itself.

If the sight of a food alone can activate the food-value system that underlies satiety, then actual eating may not be needed to initiate it. Mental engagement with food, without any eating, can indeed change appetite. Repeatedly imagining eating a food reduces its later consumption, an effect attributed to habituation of the motivational response (Morewedge et al., 2010). More broadly, observing or imagining another’s sensory or affective state can re-activate the neural circuitry that supports experiencing it first-hand (Wicker et al., 2003; Jabbi et al., 2008; Keysers & Gazzola, 2009). This evidence leaves two questions open. First, these effects required deliberate, repeated simulation, so it remains unclear whether simply watching a short video of food being eaten can change appetite. Second, this literature treats vicarious eating as a single act and does not address who is doing the eating, a distinction central to how social context shapes appetite.

Animal work shows that food value itself is malleable. In rodents, the value of a food is not fixed but is modulated by experience: mice adjust how much of an ordinary diet they eat according to whether a more- or less-preferred food is available around it, a cognitive, experience-dependent change in food value (Cheng et al., 2024). This experience-dependent malleability of food value provides an animal foundation for our approach in humans, where we ask whether value can also be updated by watching eating. We predicted that watching eating should induce satiety when the eating is one’s own but not when it is another’s, and that the two conditions should be neurally dissociable.

The more fundamental question is who is doing the eating. An identical eating event can be seen as the viewer’s own (a self-referential act) or as another’s (a third-person act), and these two interpretations recruit partly distinct systems. Eating viewed as one’s own should engage food-value and reward circuitry and induce satiety; value computed for oneself versus another is organized along the medial prefrontal cortex (Denny et al., 2012; Nicolle et al., 2012; Sul et al., 2015). The same act seen as another’s should instead engage social-cognitive circuitry and leave appetite unchanged, or even heightened, because representing another’s actions and mental states engages the mentalizing network, centered on the temporoparietal junction and dorsomedial prefrontal cortex (Saxe & Kanwisher, 2003; Van Overwalle, 2009; Schurz et al., 2014). Social context shapes appetite and food choice (Herman et al., 2003; Cruwys et al., 2015; Higgs & Thomas, 2016): the sight of food being eaten can sharpen appetite (‘visual hunger’; Spence et al., 2016), and observing another’s reward can itself be rewarding (Mobbs et al., 2009; Lockwood et al., 2015). It remains untested in humans whether the perspective taken on a single eating act (seeing the same food eaten as one’s own versus another’s, with the food and action held fixed) can dissociate appetite from its neural basis. We call satiety induced merely by watching eating, without any intake, observational sensory satiety (observational satiety for brevity); because it arises from watching rather than from eating, it can occur whether one watches oneself or another eat, and we ask whether it, and its neural basis, depend on who is seen eating.

Here we developed a video-based, individualized fMRI paradigm to test these ideas and translate the animal model to humans. Participants first rated 15 snack foods for liking and current wanting (their momentary, food-specific desire to eat each, as distinct from overall hunger or satiety) in the scanner. The four foods they rated highest for wanting were then shown as videos of the food being eaten, each either upright (a first-person view of oneself eating, self) or vertically inverted (perceived as another person eating, other). Because the two conditions are one and the same footage, the design primarily manipulates whether the eating is seen as one’s own or another’s while holding visual content, action and pre-video preference essentially constant (see Methods). We tested three hypotheses. (i) Watching oneself eat would induce greater satiety than watching another eat. (ii) Self-eating would induce satiety and a sensory-specific decrease in wanting, whereas other-eating would not. (iii) Self-eating would mainly engage food-value and reward circuitry, and other-eating mainly social-cognitive (mentalizing) circuitry. A preliminary online survey (Study 1) came first, confirming the paradigm in a demographically diverse sample and testing whether the effect holds across a broad age range, which the fMRI sample cannot. The fMRI study (Study 2) is the core of this work.

## 2. Materials and Methods

### 2.1. Stimuli

Fifteen snack foods served as the food set (Figure S1). These were everyday snack foods spanning salty and sweet types; each participant’s four experimental foods were then chosen individually as their most-wanted items (below), so the effect does not depend on the particular set. Each food was presented in two forms: a color photograph, used for the liking and wanting ratings, and a 40-s overhead video of the food being eaten, used for the self/other manipulation. In the video, a hand repeatedly picked up pieces of the snack from a plate and carried them off frame, so that the plate emptied over the 40 s. This set an unhurried but steady pace, as when eating absent-mindedly. In the self version the hand reached the food from the viewer’s own side, giving a first-person perspective of eating. The identical video was vertically inverted to create the other version, in which the hand reached the food from the opposite side, so that the video was spontaneously perceived as another person eating while facing the viewer. This manipulates viewpoint (self versus other) while holding visual content and food identity constant: the two versions are one and the same footage, vertically flipped (Figure 1). The labels ‘self’ and ‘other’ are shorthand for these first-person-upright and inverted-perspective conditions rather than a claim of verified self-experience, and a post-session questionnaire confirmed the intended perception (see below). Each video also carried the snack’s natural handling sounds. In all videos the eating hand was a single assistant’s (a 22-year-old man), not the participant’s own; the self condition was defined by the first-person viewpoint and cue, not by whose hand appeared. Each video carried a brief on-screen instruction (“Imagine that you are eating” for self, or “Imagine that another person is eating” for other). Because the footage was identical and real, this instruction served as a perspective cue rather than a request to generate a mental image with no stimulus present, directing whether the observed eating was seen as one’s own or another’s. After the session, a questionnaire confirmed that participants had perceived the inverted videos as another person, not themselves, eating.

**Figure 1.**
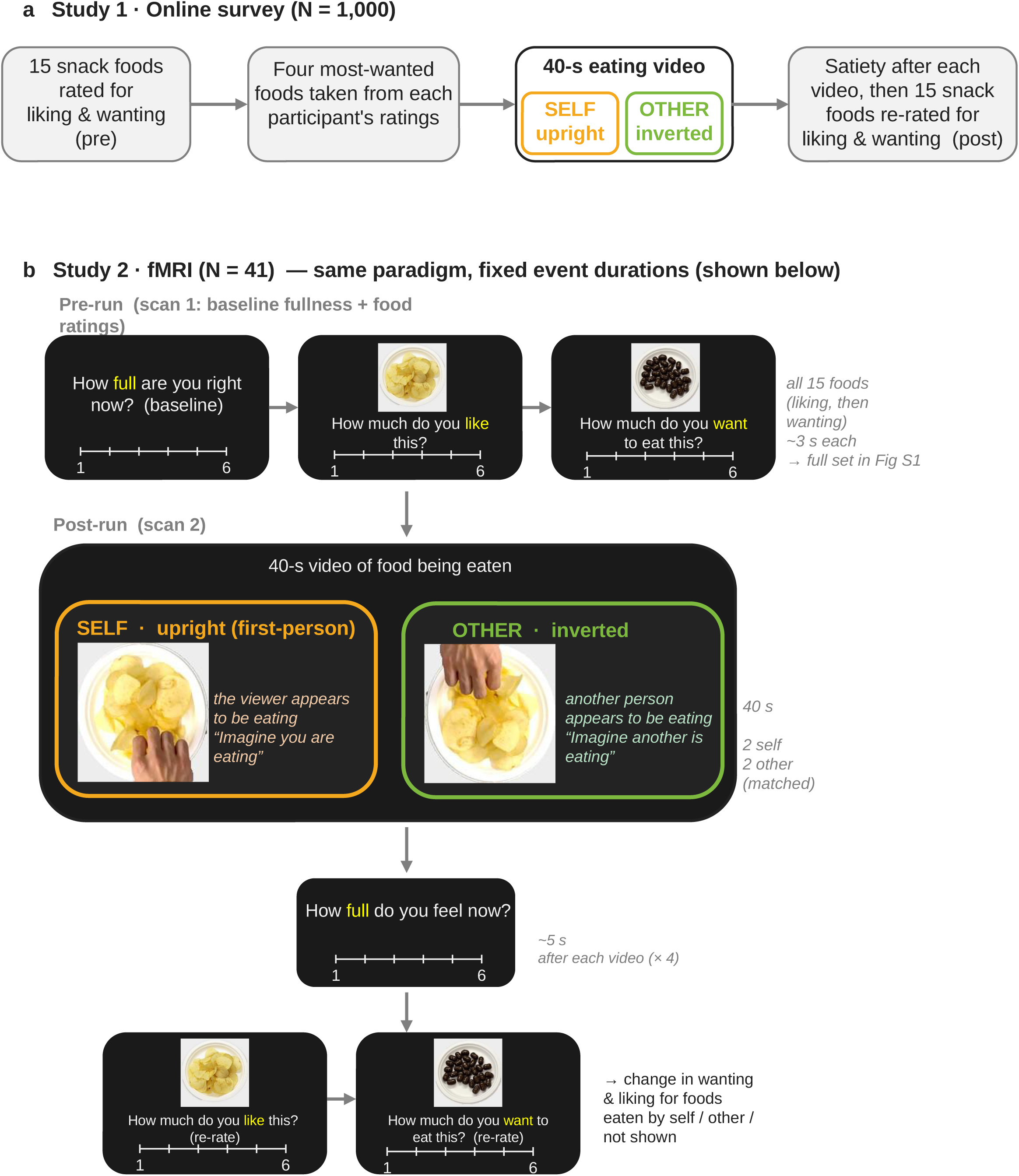
Experimental design. (a) Study 1 (online survey, N = 1,000). Participants were screened for hunger and rated all 15 snack foods for liking and wanting (6-point scales); the four foods rated highest for wanting were then presented as 40-s videos of the food being eaten. Each video was shown either upright, giving a first-person view in which the viewer appears to be eating (self), or vertically inverted, so that another person appears to be eating while facing the viewer (other); the viewpoint differs while visual content and food identity are held constant, and pre-video wanting was matched across conditions. Satiety was rated after each video, all 15 foods were re-rated to quantify the change in liking and wanting for foods eaten by the self, eaten by the other, or not shown, and two final items asked whether the videos had made participants feel full (self) or hungry (other). (b) Study 2 (fMRI, N = 41). The same paradigm implemented as timed pre- and post-video rating runs; event durations are shown (videos, 40 s; liking/wanting ratings, ∼3 s; baseline-fullness, satiety and post-video questions, ∼5 s). The full 15-food stimulus set is shown in Figure S1.

### 2.2. Study 1: online survey

We conducted an online survey of 1,000 community participants (591 men, 407 women, 2 other; mean age 52.1 ± 17.7 years, range 15–97). They were recruited through the online panel of a market-research company (Cross Marketing Inc., Tokyo, Japan). Participants were screened to have last eaten at least 2 h earlier and to feel hungry. They rated the 15 snack foods for liking and for wanting (6-point scales; 1 = not at all, 6 = extremely), and the four foods rated highest for wanting were randomly divided between self- and other-eating videos (Figure 1a). After each video they rated satiety and re-rated all 15 foods for liking and wanting. The survey placed no time limit on any response. Two further items (6-point scales) asked whether watching oneself eat a food made participants feel full, and whether watching another eat made them feel hungry, both without any actual eating. These two items indexed a general appetite state (overall fullness or hunger) rather than the food-specific wanting ratings.

### 2.3. Study 2 (fMRI): Participants

Forty-one healthy volunteers (19 men, 22 women; mean age 20.9 ± 1.2 years, range 18– 23) participated in the fMRI study. All had fasted for at least 2 h before the session, so that appetite was present but not extreme, had normal or corrected-to-normal vision, were right-handed, and reported no current neurological or psychiatric disorder. Sex was self-reported; the study was not powered for sex differences, so analyses pooled across sexes (sex comparisons are exploratory). The sample size was not set by an a-priori power calculation; a sensitivity analysis indicated that N = 41 afforded 80% power (two-sided α = 0.05) to detect a within-participant effect of dz = 0.45 (90% power for dz = 0.52), and essentially full power (>0.99) for the primary satiety effect (dz = 1.18).

### 2.4. Experimental design and procedure

In the scanner, stimuli were controlled with Presentation software (Neurobehavioral Systems, Inc., Berkeley, CA, USA) and displayed on an MR-compatible LCD monitor viewed through a mirror mounted on the head coil. Participants’ responses were obtained with an MR-compatible button box.

The task comprised two functional runs (Figure 1b). In the first (pre) run, participants rated their baseline fullness (how full they felt) and then rated all 15 foods for liking and, in a separate block, for wanting. Within each block the foods appeared one at a time, each as a photograph labeled with its name, in randomized order. The four foods each participant rated highest for wanting (ties broken at random) were then randomly assigned, two to self-eating and two to other-eating videos. In the second (post) run, these four videos were presented in randomized order. Participants rated their satiety after each video. After the four videos, two further items asked whether watching oneself eat had made them feel full and whether watching another eat had made them feel hungry (the same two probes as in Study 1), and participants then re-rated all 15 foods for liking and wanting. Comparing these with the pre-run ratings gave the change in liking and wanting for foods eaten by the self, eaten by the other, or not shown.

### 2.5. Behavioral analysis

The primary measure was post-video satiety, compared between the self and other conditions with a paired-samples *t*-test (and the Wilcoxon signed-rank test as a nonparametric check). Wanting and liking were assessed by subjective 6-point ratings and are treated as proxies for the incentive and hedonic components of food reward, respectively. Liking and wanting changes (post minus pre) were compared across foods eaten by the self, eaten by the other, and not shown, using a one-way repeated-measures ANOVA with Holm–Šidák–corrected pairwise contrasts. Effect sizes are Cohen’s dz (paired) and d (one-sample); all tests were two-tailed at α = .05. The online survey (Study 1) was analyzed with the same paired-samples and one-sample (against-zero) *t*-tests. Analyses were performed in GraphPad Prism 10 (GraphPad Software, Boston, MA, USA).

### 2.6. MRI data acquisition

MRI data were acquired on a 3.0-T GE Healthcare SIGNA Premier scanner (software RX29.1) with a 48-channel head coil. Functional volumes were obtained with a T2*-weighted gradient-echo echo-planar imaging sequence. Acquisition parameters were: repetition time [TR] = 3000 ms, echo time [TE] = 30 ms, flip angle = 90°, 38 axial slices acquired in ascending order, slice thickness = 3.0 mm with no gap, field of view = 240 mm, acquisition matrix = 80 × 80 [in-plane resolution 3.0 × 3.0 mm]. Two functional runs were collected: a pre run (42 volumes) containing the baseline fullness and food-rating task, and a post run (106 volumes) containing the video-viewing and re-rating task. At the start of each run, dummy volumes were acquired and discarded over an initial 12-s period, so magnetization was already at steady state at the first stored volume, and no volumes were discarded from the analysis. A high-resolution sagittal 3D T1-weighted anatomical image (MPRAGE with prospective motion correction; TR = 2723 ms, TE = 2.0 ms, inversion time = 1000 ms, flip angle = 8°, field of view = 240 mm, 0.9-mm isotropic voxels) was acquired between the two functional runs.

### 2.7. fMRI preprocessing and analysis

Data were preprocessed and analyzed with SPM12 (Wellcome Centre for Human Neuroimaging, London, UK) running under MATLAB R2024b (The MathWorks, Natick, MA, USA). Preprocessing comprised slice-timing correction, realignment and unwarping, coregistration of the mean functional image to the segmented T1 image, spatial normalization to MNI space via the T1 forward deformation field (functional images resampled to 2 × 2 × 2 mm), and smoothing with an 8-mm full-width-at-half-maximum Gaussian kernel.

#### First-level (individual) models

All general linear models used the canonical hemodynamic response function, a 128-s high-pass filter, and an AR(1) model for serial correlations, with the six head-motion parameters as nuisance regressors. Six complementary models were specified, numbered in the order they first appear in the Results. (1) A model splitting each video into four 10-s bins examined the temporal dynamics of the self–other difference. (2) A pre/post model of liking and wanting for self-, other-, and not-shown foods isolated the sensory-specific change in wanting, free of the visual differences between the self and other videos. (3) A satiety-rating model that also modeled the button-press component contrasted self-versus other-eating during the satiety judgment within a-priori appetite regions. (4) A base categorical model of the post run coded the self- and other-eating videos (40-s epochs) and the rating events; model comparison (Bayesian Information Criterion) supported this event-based specification over parametric variants. (5) A value-parametric model of the pre-run localized activity scaling with rated wanting (or liking) during food-photo viewing, before any videos. (6) A psychophysiological-interaction model seeded on the right temporoparietal junction tested task-dependent coupling during the self versus other conditions.

#### Individual-difference analyses

To test whether appetite influenced this activity, we related a-priori ROI signals (mean β within the same 8-mm spheres) to individual differences in appetite state (baseline pre-run fullness) using Pearson and Spearman correlations across the 41 participants. These brain–behavior analyses were exploratory and are reported as such.

#### Second-level (group) analysis

First-level contrast images were carried to second-level one-sample t-tests across all 41 participants. Results are reported at a voxel-forming threshold of p < .001 (uncorrected) with cluster-level family-wise-error (FWE) correction at p < .05 for whole-brain inference (Eklund et al., 2016). For a-priori regions of interest (insula, ventral striatum, amygdala, orbitofrontal and medial prefrontal cortex, and the temporoparietal-junction/mentalizing network), small-volume FWE correction (p < .05) was applied within 8-mm-radius spheres centered on coordinates drawn from prior literature. Anatomical labels and MNI coordinates of local maxima are given in Table 1, and unthresholded statistical maps are available at https://neurovault.org/collections/24249/.

**Table 1.** Local maxima for the principal contrasts (N = 41). Peak t, cluster extent k (voxel-forming threshold p < .001 uncorrected), and corrected significance: whole-brain cluster-level p_FWE, or small-volume peak p_FWE (SVC) within an 8-mm-radius a-priori ROI.

| Region | Side | x | y | z | k | T | p | Correction |
| --- | --- | --- | --- | --- | --- | --- | --- | --- |
| **Food viewing (pre-video ratings) > baseline — insula** |  |  |  |  |  |  |  |  |
| Insula | L | -38 | -6 | 12 | 887 | 6.57 | <.001 | peak |
| Insula | R | 40 | -2 | 10 | 9 | 3.67 | .011 | SVC |
| **Video viewing, temporal accumulation (linear ramp, self + other)** |  |  |  |  |  |  |  |  |
| Ventromedial OFC / PFC | — | -4 | 50 | -10 | — | 7.11 | .001 | peak |
| Ventral striatum | R | 20 | 10 | -6 | — | 7.18 | <.001 | peak |
| Ventral striatum | L | -12 | 10 | -6 | — | 6.75 | .001 | peak |
| **Video viewing, other > self — mentalizing network** |  |  |  |  |  |  |  |  |
| TPJ / angular gyrus | R | 36 | -66 | 20 | 680 | 4.94 | .0004 | cluster |
| TPJ / angular gyrus | L | -54 | -56 | 26 | — | 3.05 | .14 (n.s.) | SVC |
| Dorsomedial PFC | — | 0 | 52 | 28 | — | 2.76 | .13 (n.s.) | SVC |
| Precuneus / PCC | — | 0 | -56 | 34 | — | 2.89 | .063 | SVC |
| Posterior STS | R | 52 | -40 | 4 | — | 2.63 | .10 (n.s.) | SVC |
| Temporal pole | R | 58 | 4 | -18 | 52 | 4.42 | .061 | SVC |
| **Video viewing, self > other — attention / visual** |  |  |  |  |  |  |  |  |
| Superior parietal lobule | L | -28 | -56 | 64 | 629 | 5.76 | n.s. | cluster |
| Middle occipital / calcarine | L | -8 | -98 | -2 | 49 | 4.50 | n.s. | cluster |
| **Sensory-specific satiety of wanting (shown vs not-shown, post-pre)** |  |  |  |  |  |  |  |  |
| TPJ / angular gyrus | L | -54 | -52 | 32 | 208 | 4.33 | .040 | cluster |
| TPJ / angular gyrus | R | 56 | -60 | 34 | 150 | 4.20 | .004 | SVC |
| Inferior frontal gyrus | R | 46 | 36 | 14 | 69 | 4.26 | .003 | SVC |
| Medial PFC | — | 12 | 54 | 24 | — | 3.98 | .061 | SVC |
| **Shown foods, post > pre** |  |  |  |  |  |  |  |  |
| Inferior parietal lobule | L | -54 | -48 | 38 | 960 | 5.54 | $8 \times 10^{-6}$ | cluster |
| Supramarginal gyrus | R | 58 | -46 | 36 | 918 | 4.95 | $1 \times 10^{-5}$ | cluster |
| Putamen | R | 28 | 14 | 0 | 751 | 5.25 | $7 \times 10^{-5}$ | cluster |
| Caudate | R | 12 | 14 | 8 | — | 4.34 | — | cluster |
| Precuneus | L | -12 | -64 | 34 | 640 | 5.17 | $2 \times 10^{-4}$ | cluster |
| Medial PFC / dACC | L | -10 | 24 | 44 | 287 | 5.23 | .016 | cluster |
| **Shown vs not-shown interaction (liking + wanting; food-specificity control)** |  |  |  |  |  |  |  |  |
| Ventral striatum | L | -6 | 12 | -4 | — | 3.29 | .034 | SVC |
| Insula | R | 36 | 12 | -2 | — | 3.58 | .023 | SVC |
| TPJ / angular gyrus | L | -48 | -56 | 34 | — | 3.04 | .049 | SVC |
| Medial PFC | — | 0 | 50 | 12 | — | 1.92 | .36 (n.s.) | SVC |
| **Wanting change, self > other** |  |  |  |  |  |  |  |  |
| Mid / dorsomedial PFC | L | -10 | 46 | 22 | 28 | 4.22 | .004 | SVC |
| Ventral striatum / putamen | L | -12 | 10 | -6 | — | 3.39 | .026 | SVC |
| Ventral striatum / putamen | R | 12 | 10 | -6 | — | 2.34 | .18 (n.s.) | SVC |
| **Satiety judgment, self > other (motor-controlled; a-priori appetite ROIs — none significant)** |  |  |  |  |  |  |  |  |
| Amygdala | L | -24 | -4 | -18 | — | 1.71 | .34 (n.s.) | SVC |
| Amygdala | R | 24 | -4 | -18 | — | 1.70 | .35 (n.s.) | SVC |
| Ventral striatum | R | 12 | 10 | -6 | — | 0.95 | .56 (n.s.) | SVC |
| Insula | R | 38 | 6 | 2 | — | 0.88 | .58 (n.s.) | SVC |
| Medial PFC | — | 0 | 50 | 12 | — | 0.83 | .59 (n.s.) | SVC |
| Orbitofrontal cortex | — | 0 | 44 | -12 | — | 0.45 | .65 (n.s.) | SVC |
| **Wanting-parametric, food-photo viewing (a-priori ROI mean $\beta$, one-sample t; liking-parametric: all n.s.)** |  |  |  |  |  |  |  |  |
| Orbitofrontal cortex | — | 0 | 44 | -12 | — | 4.01 | $3 \times 10^{-4}$ | ROI t |
| Medial PFC | — | 0 | 50 | 12 | — | 2.64 | .012 | ROI t |
| Amygdala | R | 24 | -4 | -18 | — | 2.07 | .045 | ROI t |
| Ventral striatum | R | 12 | 10 | -6 | — | 1.75 | .088<br>(n.s.) | ROI t |
| Insula | R | 38 | 6 | 2 | — | 1.42 | .16 (n.s.) | ROI t |

Coordinates are peak MNI coordinates (mm); k, cluster extent in voxels (voxel-forming threshold $p < .001$ uncorrected); T, peak t; p, corrected significance obtained with the procedure named in the Correction column. Correction: peak, whole-brain peak-level FWE (sign-flip max-t permutation, 2000 iterations); cluster, whole-brain cluster-level FWE; SVC, small-volume-corrected peak FWE within an 8-mm-radius a-priori ROI; ROI t, one-sample-t p across participants for the a-priori ROI mean parametric slope ( $\beta$ ). Rows marked n.s. under cluster correction had no cluster survive whole-brain FWE. The food-viewing contrast is from a pre-run food-cue model (all food photos versus baseline, before any video); the sensory-specific-satiety and post > pre contrasts are from the viewpoint-matched pre/post model; the temporal-accumulation slopes are from the 10-s-bin model and were steeper than in the visual control region V1 (OFC > V1, paired $t(40) = 4.70$ , $p < .0001$ ); the video other > self and self > other contrasts are from the base categorical model.

## 3. Results

### 3.1. Preliminary survey (Study 1): the behavioral effect holds across a broad age range

Before the fMRI study, an online survey supported the paradigm and tested its generality; the 15-food set was well balanced across participants (Figure S1). Satiety was higher after watching oneself eat than after watching another eat (*p* = 2.5 × 10⁻⁹; Figure 2a). Wanting for the eaten foods decreased and wanting for the not-shown foods increased, whereas liking changed little in any condition and showed no such food-specific pattern (Figure 2b). This raw shown-versus-not-shown contrast in wanting is, however, inflated by regression to the mean, because each participant’s eaten foods were their most-wanted and so started near the top of the scale: when eaten and not-shown foods were matched within participants on both pre-video wanting and liking, the difference became negligible (Δ ≈ +0.01, n.s.). (Operationally, within each participant we paired eaten and not-shown foods to have matched pre-video wanting and liking, then compared the wanting change between these matched sets.) The wanting decrease was, moreover, equal for self- and other-eaten foods (*p* = 1.0). The reported satiety, by contrast, was self-specific and cannot be explained by regression to the mean, because the self- and other-eaten foods were drawn from the same most-wanted set and differed only in viewpoint; it is this attribution-dependent satiety that the fMRI study was designed to localize. Importantly, the effects generalized across sex and a broad cross-sectional age range (sex, both *p* > .16; age, *r* = .03, *p* = .32; Figure S2a,b). The decrease also did not depend on taste (salty versus sweet) or color (yellow/white versus brown; Figure S2c). After the videos, two further questionnaire items probed a possible ‘vicarious hunger’. One asked how much watching oneself eat had made participants feel full, and the other how much watching another eat had made them feel hungry. Participants agreed slightly more with the hunger item than with the fullness item, although both ratings stayed below the scale midpoint (Figure 2c).

**Figure 2.**
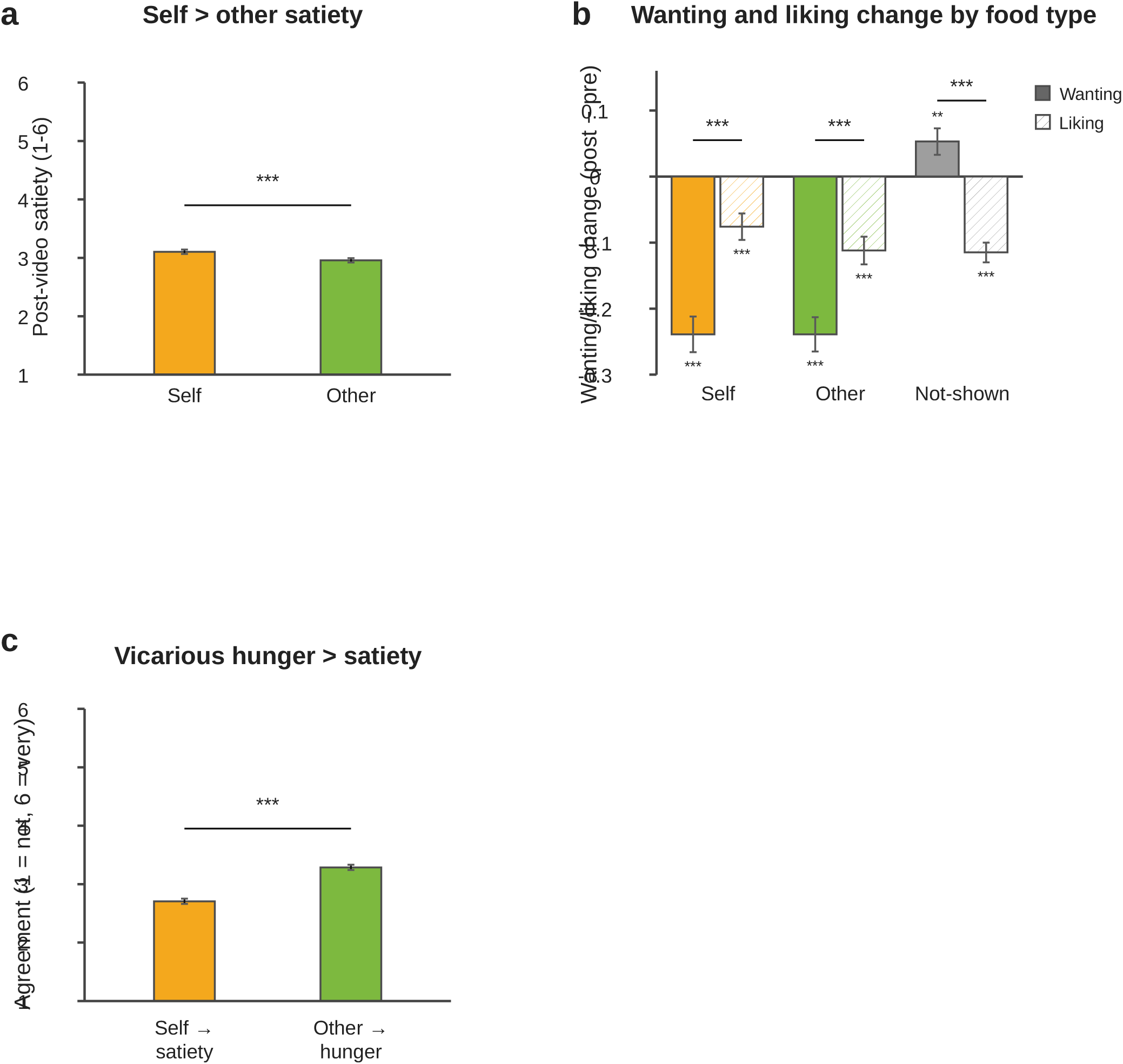
Large-scale online survey (Study 1; N = 1,000; ages 15–97). (a) Satiety, rated on a 6-point scale (1 = not at all, 6 = very much), after watching oneself eat versus watching another eat. (b) Wanting and liking change (post − pre), each split by condition (self-eaten, other-eaten, not-shown); brackets compare wanting versus liking within each condition, and stars below the bars test each change against zero. (c) Two self-report items: agreement that watching another person eat increased one’s own hunger, and that watching oneself eat induced satiety (6-point agreement scale, 1 = not at all, 6 = very much). Bars, mean; whiskers, s.e.m.

### 3.2. The fMRI study (Study 2): watching oneself eat induces satiety without eating

Within each participant, the foods shown in the self and other videos had been wanted equally before the videos, so any later difference in satiety reflects the change in perspective rather than a difference between the foods. As in Study 1, satiety after the videos was higher in the self than in the other condition (paired *t*(40) = 7.54, *p* = 3.4 × 10⁻⁹, Cohen’s *d_z_* = 1.18, 95% CI [0.84, 1.69]; Figure 3a), showing that a brief video induced satiety (an observational sensory satiety, that is, satiety induced by watching eating rather than by eating itself) only when the eating was seen as one’s own.

**Figure 3.**
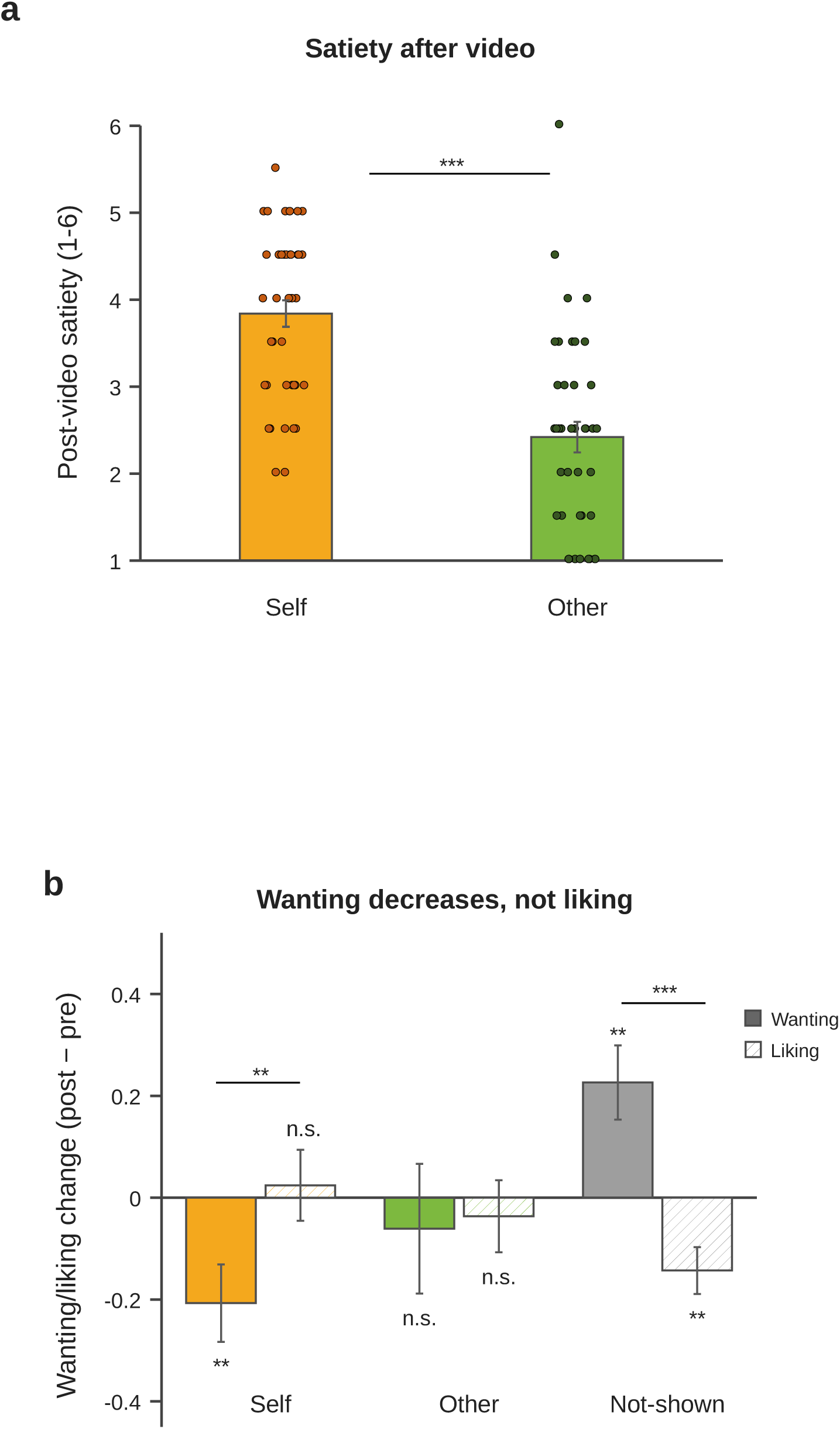
Behavioral results (N = 41). Reported satiety was greater for self than other, whereas the food-specific decrease in wanting did not differ significantly between self- and other-eaten foods, though it was weaker and more variable for other-eaten foods (Figure S3): a partial dissociation between subjective satiety and food devaluation. (a) Satiety (6-point scale, 1 = not at all, 6 = very much) after self-versus other-eating videos (bars, mean ± s.e.m.; gray points, individual participants). (b) Wanting (solid) and liking (hatched) change (post − pre) for foods eaten by self, eaten by other, and not shown; brackets compare wanting versus liking within each condition, and stars below the bars test each change against zero. Bars, mean; whiskers, s.e.m. \*\*\**p* < .001, \*\**p* < .01, \**p* < .05; n.s., not significant.

The post-video re-ratings showed a selective, food-specific decrease in wanting but not liking. Wanting change differed across the three food types (self-eaten, other-eaten and not-shown; one-way repeated-measures ANOVA, *F*(2,80) = 8.01, *p* = 6.7 × 10⁻⁴): it decreased for the foods shown being eaten and increased for the not-shown foods. The decrease was significant for self-eaten foods (versus not-shown, *t*(40) = −5.75, *p* = 3.2 × 10⁻⁶, *d_z_* = −0.90; Figure 3b) and weaker, and only relative to the not-shown foods, for other-eaten foods (*t*(40) = −2.36, *p* = .046). In this smaller sample the self- and other-eaten changes did not differ statistically (*p* = .25). Unlike in the large online survey (Study 1), the other-eaten change did not itself fall below baseline in this smaller sample. The other condition, however, was more variable across participants, with wanting decreasing in some and increasing in others (Levene’s test, *p* = .026; Figure S3). Liking change, by contrast, was small and did not differ across the three food types (*F*(2,80) = 2.33, *p* = 0.10; Figure 3b). As in Study 1, the decrease in wanting was not moderated by sex or age (all *p* > .2) and did not differ by taste (salty versus sweet, *p* = .51) or color (yellow/white versus brown; Figure S2d).

### 3.3. While watching, a food-value signal common to self and other

In the brain, the two conditions first shared a common signal. Even before any video was shown, viewing the food photos during the pre-video ratings engaged the insula, a taste- and interoceptive region that responds to the mere sight of food (Small, 2010; van der Laan et al., 2011), bilaterally (left insula whole-brain FWE, peak *t* = 6.57; right insula small-volume-corrected, *t* = 3.67, *p*_FWE_ = .011; Figure 4a). During the videos, the a-priori appetite regions (insula, ventral striatum, amygdala, orbitofrontal and medial prefrontal cortex) showed a common food-related signal. Their BOLD signal increased steadily across the 40-s video and did not differ between watching oneself and watching another in any region (Figure S4; linear ramp all *p* < .01 except the insula, *p* = .056; self versus other n.s.). This build-up was largest in the ventromedial orbitofrontal cortex and ventral striatum, which increased more steeply than the visual control region V1 (10-s-bin model, model 1; OFC > V1 ramp, *p* < .0001; ventral striatum whole-brain FWE; Figure 4b). As a negative control, V1 itself showed no such ramp, so the build-up was specific to value regions rather than a generic visual or signal-drift effect. Viewing food being eaten thus engaged food-value circuitry irrespective of whose eating it was.

**Figure 4.**
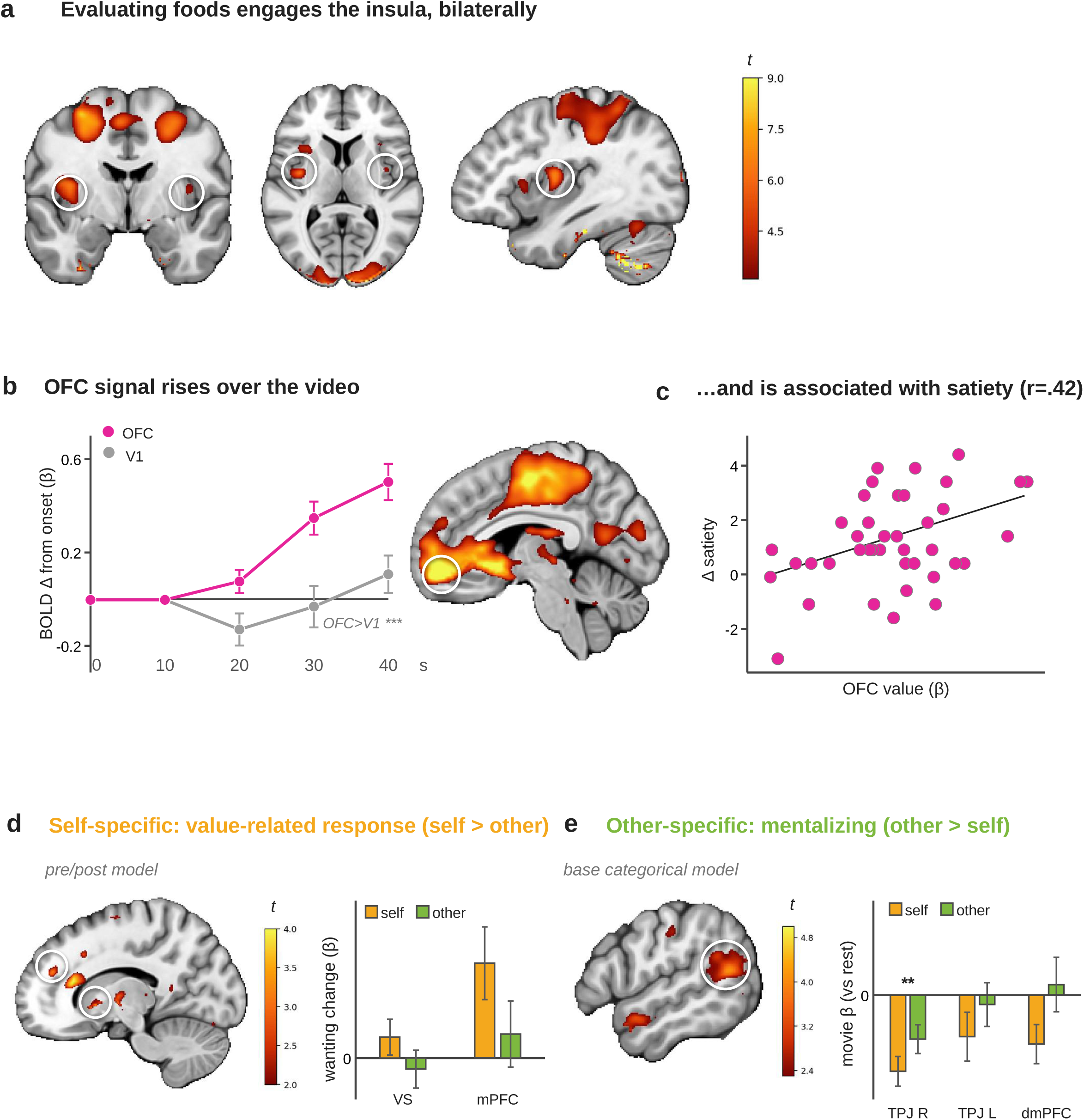
From food evaluation to attribution-specific appetite change (N = 41). (a) Pre-video food evaluation. During the pre-video food ratings, before any video was shown, the bilateral insula was engaged (left insula whole-brain FWE, right insula small-volume FWE); the dorsal midline cluster reflects the rating button-press. (b) In a 10-s-bin model, ventromedial orbitofrontal cortex and ventral striatum activation across the 40-s video, shown against the visual control region V1 (OFC > V1 ramp, *p* < .0001), for watching oneself and watching another (curves, change from video onset; ventral striatum whole-brain FWE); the common, all-ROI form is shown in Figure S4. (c, exploratory) Ventromedial-OFC activation while watching oneself eat plotted against each participant’s satiety increase (*r* = .42, *p* = .006); the circled OFC in (b) is the source region for (b) and (c). (d) The self > other contrast of the wanting change (pre/post model): left ventral striatum and medial prefrontal cortex (small-volume-corrected, *p* < .05). (e) Watching another eat (other > self, base categorical model): right temporoparietal junction (whole-brain cluster-FWE, *p* = .0004) and dorsomedial prefrontal cortex. Brain slices, group T-maps on white-cut-out anatomical (sagittal shown anterior-left); color bars, t. See Table 1 for local maxima and corrected p-values.

### 3.4. Watching food being eaten leaves a change specific to the eaten foods

Beyond this common signal, watching food being eaten changed how the brain represented the shown foods. Using the viewpoint-matched pre/post model (model 2), we compared the response to the shown (eaten) foods, as participants re-rated them for liking and wanting, after versus before the videos, with the same food photographs shown before and after each video, so that only the intervening video differed. Relative to the not-shown foods, this pre-to-post change was specific to the eaten foods. In the wanting-specific contrast (wanting change, shown versus not-shown), the left temporoparietal junction/angular gyrus survived whole-brain correction (*k* = 208, *p*_FWE_ = 0.040; bilateral TPJ small-volume *p*_FWE_ = 0.003/0.004), with no liking counterpart, matching the wanting-specific behavior. A categorical shown-versus-not-shown interaction (pooling liking and wanting) survived small-volume correction in the left ventral striatum (peak *t* = 3.29 at [−6, 12, −4], *p*_FWE_ = .034), the right insula ([36, 12, −2], *p*_FWE_ = .023) and the left temporoparietal junction (*p*_FWE_ = .049), but not in the medial prefrontal or orbitofrontal cortex, and no region survived whole-brain voxel-level correction (Table 1). The ventral-striatal and insular value-related responses were therefore specific to the shown foods; a medial-prefrontal cluster seen in the simpler post > pre contrast did not survive this shown-versus-not-shown control (it appeared for the not-shown foods as well) and coincided with a general pre-to-post signal decrease even in the visual control region (*t*(40) = −2.82, *p* = .007), so we treat the medial-prefrontal change as non-specific. When instead time-locked to the satiety rating itself (motor-controlled model 3), a-priori taste and food-value regions (insula, orbitofrontal cortex) did not reach significance (all small-volume *p*_FWE_ > 0.3). The change was thus carried by food value and wanting for the shown foods, not by a primary gustatory signal.

### 3.5. A self-specific value-related response that scales with reported satiety

Two findings tied a value-related response specifically to eating seen as one’s own and to the satiety it produced. First, orbitofrontal activation while watching oneself eat scaled with the satiety each participant gained. The more a participant’s orbitofrontal cortex responded during the self-eating videos, the larger their increase in satiety (*r* = .42, *p* = .006; Spearman ρ = .31; Figure 4c). Second, the self > other contrast of the wanting change engaged the mid-/dorsomedial prefrontal cortex (*t* = 4.22, small-volume-corrected *p*_FWE_ = 0.004) and the left ventral striatum/putamen (*p*_FWE_ = 0.026; Figure 4d), although at the a-priori ROI level these self-versus-other differences (self-eaten minus other-eaten wanting-change activation, averaged within each 8-mm ROI) were only trend-level. These brain– behavior relationships are exploratory and uncorrected, but together they point to the same conclusion. Greater engagement of food-value regions while watching themselves eat was associated with more reported satiety.

### 3.6. The same act seen as another’s engages a mentalizing region instead

The same act seen as another’s recruited a different system. Inverting the video made the identical, equally-wanted food appear to be eaten by someone else, and this viewpoint manipulation was neurally effective. In the base categorical model (model 4), watching another eat (other > self) engaged the right temporoparietal junction/angular gyrus, which survived whole-brain cluster-level correction (*k* = 680, *p*_FWE_ = 0.0004; small-volume *p*_FWE_ = 0.003). Sub-threshold activation appeared in the left TPJ, dorsomedial prefrontal cortex, precuneus, right posterior superior temporal sulcus and temporal pole (Figure 4e; the mentalizing-ROI summary is shown in Figure S5). Activation extracted directly from the right-TPJ region showed a reliable other > self difference across the group (mean β = +0.15, *t*(40) = 3.08, *p* = .004). The reverse contrast (self > other; that is, regions more active when watching oneself eat than another) activated only the left superior parietal lobule (*t* = 5.76) and occipital cortex, a dorsal-attention and visual pattern attributable to the low-level difference between the upright and inverted videos rather than to perspective. Unlike the self-specific value-related response, the magnitude of this other-specific temporoparietal activation did not scale with any appetite measure across participants, including the survey’s possible ‘vicarious hunger’ (all |*r*| < .31, *p* > .05). The region was recruited by another’s eating, but individual differences in its activation did not predict individual appetite change. Seeing the act as another’s thus recruited a mentalizing region, the social-cognitive network for representing another’s actions and mental states, whereas the value-related response and reported satiety remained specific to eating seen as one’s own. This is the neural dissociation predicted in (iii).

Underlying these effects (the video-evoked value and mentalizing responses described above), the valuation regions already coded food value before any video was shown. In the pre-run value-parametric model (model 5), activity during food-photo viewing scaled with each food’s rated wanting in the orbitofrontal cortex (a-priori ROI, *t*(40) = 4.01, *p* = 3 × 10⁻⁴, *d_z_* = 0.63; small-volume-corrected peak *t* = 5.01, *p*_FWE_ < .05), the medial prefrontal cortex (*t*(40) = 2.64, *p* = .012) and the amygdala (*t*(40) = 2.07, *p* = .045), with a comparable trend in the ventral striatum (*t*(40) = 1.75, *p* = .088; Figure 5). No region scaled with liking (all *p* > .19), the same wanting-specific pattern found in the behavior. These value signals also did not differ between salty and sweet or yellow/white and brown foods (all *p* > .17; Figure S2e,f).

**Figure 5.**
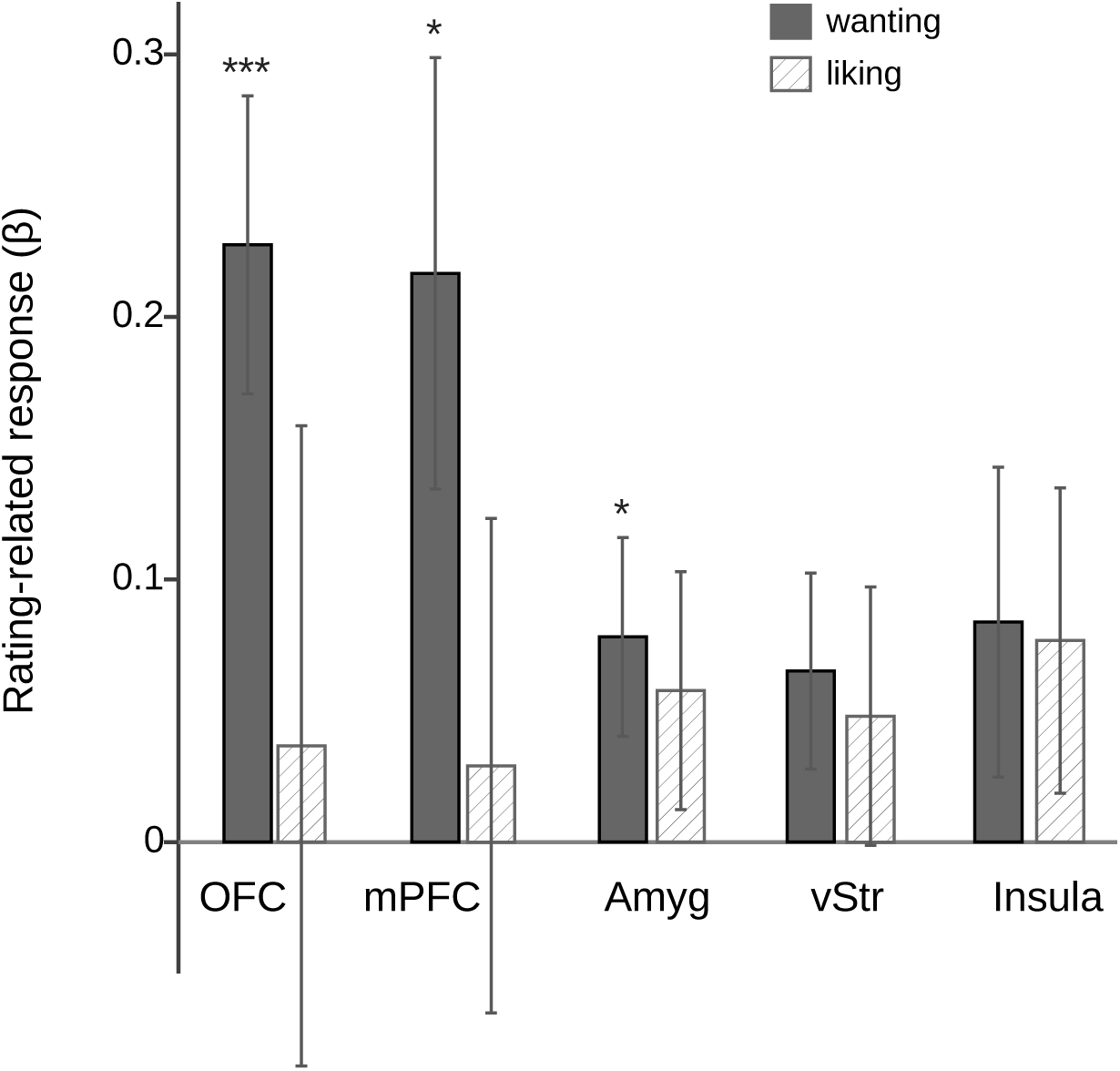
Food value is coded by wanting (N = 41). During food-photo viewing, activity scaled with rated wanting in the orbitofrontal and medial prefrontal cortex and amygdala, but not with liking; a visual control region showed no effect. Wanting is shown as solid gray bars and liking as hatched bars (diagonal fill), matching the convention in Figures 2b and 3b. Bars, mean ± s.e.m.; points, participants (small vertical jitter on the discrete rating scales). *p < .05, **p < .01, ***p < .001.

### 3.7. Specificity: model-comparison and connectivity controls

Two control analyses confirmed that these conclusions did not depend on modeling choices or reflect region-to-region connectivity (Supplementary Methods and Results). These compared alternative ways of modeling the eating videos and tested directed connectivity between the mentalizing and value regions.

## 4. Discussion

Watching oneself eat induced satiety without any eating. Our three a-priori predictions were largely supported. First, self-eating induced greater satiety than other-eating within the same, matched participants (predictions i and ii). Second, this satiety was accompanied by a food-specific decrease in wanting for the eaten foods. This decrease was clearest for self-eaten foods; in the large online survey (Study 1) it extended to other-eaten foods as well, though in the smaller fMRI sample the other-eaten decrease was weak and variable: observing a food being eaten can devalue it largely regardless of agent, and only seeing the eating as one’s own converts this devaluation into reported satiety (a refinement of prediction ii). Third, the two conditions were neurally dissociable: eating seen as one’s own engaged reward and valuation circuitry, whereas eating seen as another’s engaged the social-cognitive mentalizing network (prediction iii). The same eating act was associated with a self-specific food-value response and greater reported satiety when seen as one’s own, but engaged the social-cognitive system and less satiety when seen as another’s (Figure 6).

**Figure 6.**
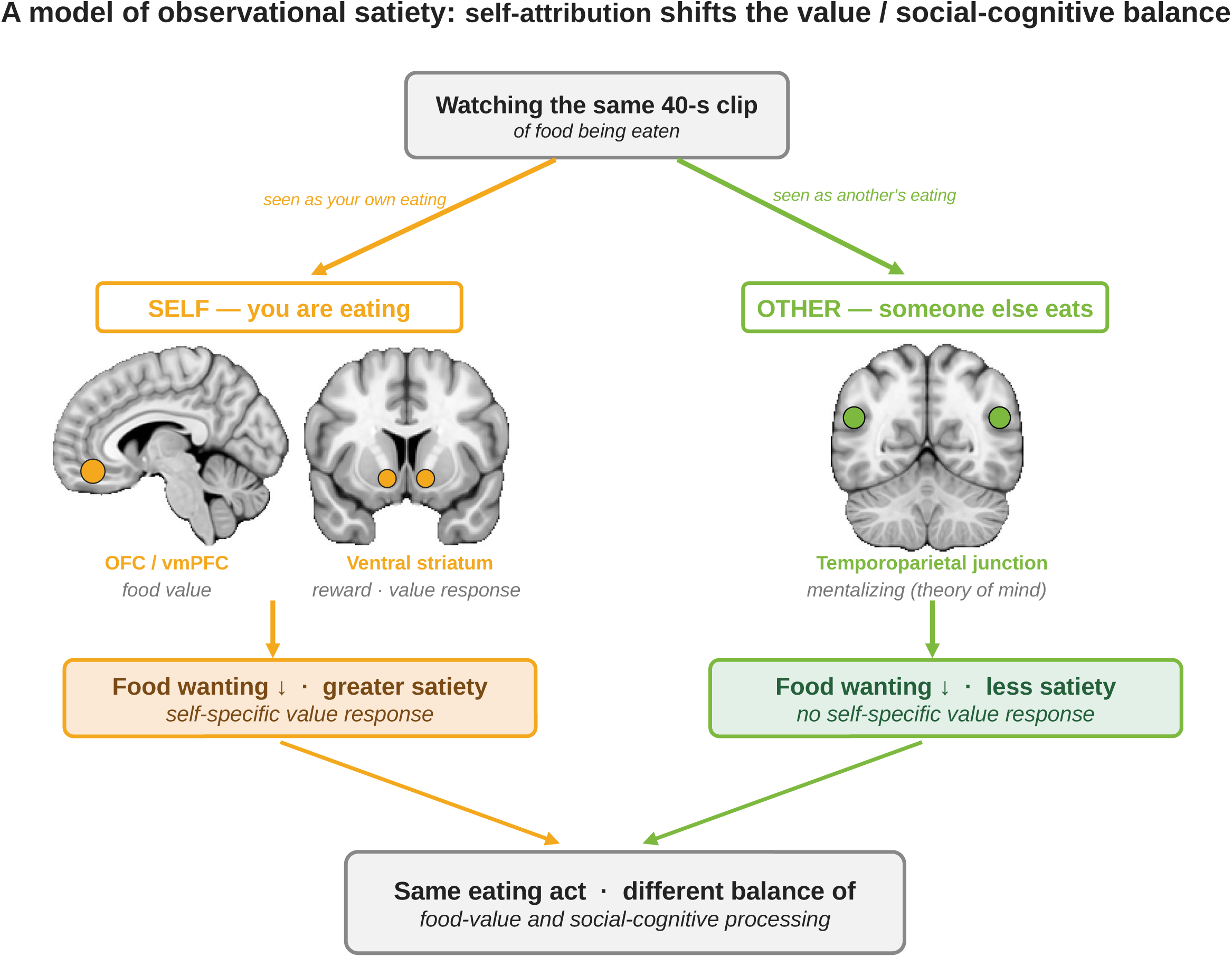
Proposed model: self-attribution determines observational satiety. Watching an identical 40-s video of food being eaten changes appetite in opposite directions depending on whether the eating is seen as one’s own. When the act is seen as one’s own, reward and valuation circuitry (ventral striatum, medial prefrontal cortex) shows a self-specific value-related response; wanting for the eaten food selectively decreases and more satiety is reported, without any eating. When the identical act is seen as another’s, the social-cognitive (mentalizing) system (temporoparietal junction, dorsomedial prefrontal cortex) is engaged instead, with no self-specific value-related change and less reported satiety. A large online survey (N = 1,000; ages 15–97) showed that the behavioral effect holds across a broad age range. The figure summarizes the account developed in the Discussion and may also serve as a graphical abstract.

### 4.1. Observational satiety reflects a change in food value

The satiety induced by self-eating was expressed as a change in food value rather than as a primary gustatory signal. A self > other contrast of the wanting-related activation engaged the ventral striatum and medial prefrontal cortex at the small-volume-corrected level, even though the behavioral decrease in wanting itself did not differ between self- and other-eaten foods, and orbitofrontal activation while watching oneself eat scaled with the satiety each participant gained. A shown-versus-not-shown interaction confirmed that this pre-to-post change was specific to the eaten foods in the left ventral striatum, right insula and left temporoparietal junction (small-volume-corrected), though not in the medial prefrontal cortex; the medial-prefrontal cluster seen in the simpler post > pre contrast did not survive this control and is treated as non-specific. By contrast, a-priori taste and food-value regions (insula, orbitofrontal cortex) were not significantly engaged during the satiety rating itself. This suggests that a purely observational video updates the motivational value of food without fully re-creating a first-hand taste or bodily experience. This fits the finding that imagined consumption can reduce subsequent eating, an effect attributed to habituation of the motivational (wanting) response rather than to a change in liking (Morewedge et al., 2010). Because our effect likewise appeared in wanting but not in liking, it corresponds to this motivational, food-specific form of devaluation rather than to classical sensory-specific satiety, which is indexed mainly by a decline in pleasantness (E. T. Rolls, 2015; Finlayson et al., 2007). We therefore reserve the term satiety for the subjective fullness that participants reported, which was greater when the eating was seen as one’s own. This also differs from work linking the brain’s baseline responsiveness to food reward to how much people eat (Adise et al., 2018): rather than relating reward sensitivity to intake, our results show that observing a food being eaten changes its value, and that whether this devaluation is accompanied by satiety depends on self-attribution. Because participants never tasted the food, hedonic (liking) updating, which may require real sensory exposure, was not engaged, whereas the food’s motivational value (wanting) was revised from observation alone. A fuller taste or bodily signature might emerge with greater statistical power or a more immersive manipulation.

This wanting-specific satiety (satiety accompanied by a food-specific fall in wanting) is not unique to the vicarious paradigm. In a separate experiment, six participants who actually ate a food for five minutes showed the same taste-graded decrease in wanting, but not liking. This provides preliminary convergent evidence that the video effect reflects a sensory-specific-satiety process rather than only a report bias (Supplementary Methods and Results; Figure S6).

### 4.2. Eating seen as another’s engages the mentalizing network

Watching the same, equally-wanted food eaten by another induced less satiety. Instead it recruited the mentalizing network that supports representing others’ actions and mental states (Saxe & Kanwisher, 2003; Van Overwalle, 2009): the bilateral temporoparietal junction, dorsomedial prefrontal cortex, precuneus, posterior superior temporal sulcus and temporal pole. The right temporoparietal junction is also engaged by attentional reorienting, so this cluster on its own would be ambiguous; its co-activation with the rest of this distributed network, and a peak dorsal and posterior to the ventral attention focus, favors a mentalizing rather than an attentional interpretation. Because the self and other videos depicted the identical eating action and differed only in viewpoint, any action-observation (‘mirror’) activation, which would engage ventral premotor and inferior parietal cortex, was common to both conditions and canceled out in their contrast. The residual other > self difference therefore indexes whether the eating was seen as one’s own rather than the movement itself, consistent with the distinction between the mirror and mentalizing systems (Van Overwalle, 2009). The same regions showed the food-specific decrease in wanting in the viewpoint-matched pre/post analysis, linking the social interpretation of eating to the update of the viewer’s own food value. One interpretation is that, seen as another’s, the eating is mentally represented as someone else’s act rather than vicariously “consumed” as one’s own. Wanting for the eaten food nonetheless decreased for self-eaten foods and, most clearly in the large survey, for other-eaten foods too, consistent with observed eating devaluing a food to some degree whichever agent eats it. These findings place food-related self–other processing within domain-general social-cognitive circuitry.

This dissociation is grounded first in behavior, and only then in the neural data. The two conditions showed the identical eating action and differed only in whether it was seen as one’s own or another’s, so any action-observation or ‘mirror’ resonance was matched across them. On a simple shared-circuits account, in which watching eating re-activates the observer’s own consummatory representation, observing the action should sate the viewer whoever is seen eating. Behaviorally it did not: fullness rose after watching one’s own eating but not after watching another’s (a self-specific increase, dz = 1.18), even though the observed action was the same. What separated the two conditions was the attribution of the eating to the self, not the movement. The decrease in wanting was less agent-specific, consistent with observed eating devaluing a food to some degree whichever agent eats it, which places the agency effect at whether this devaluation is registered as the viewer’s own satiety rather than at observation itself. The neural data then supply a mechanism consistent with this behavioral pattern: the food-value representation was shared across self and other, whereas watching another additionally engaged the mentalizing network that represents others’ actions and mental states, rather than the ventral premotor and inferior parietal mirror system. A self-versus-other attribution, computed by the mentalizing rather than the mirror system, thus gates whether a shared representation of eating is converted into a change in the viewer’s own appetite, a principle that need not be limited to eating.

A further possibility is a potential mirror image of the observational satiety reported here: watching another eat may not merely fail to satiate but somewhat raise the viewer’s appetite, a candidate ‘vicarious hunger’ (an observation-induced, sensory-specific hunger) that fits the ‘visual hunger’ literature and the reward responses to others’ eating (Spence et al., 2016; Järvinen et al., 2025). Our data provide only indirect evidence. The vicarious-hunger self-report was endorsed more than the observational-satiety one, though both stayed below the scale midpoint and other > self wanting was absent. The wanting response to watching another eat was also heterogeneous: some viewers devalued the food, whereas others wanted it more (Figure S3). If confirmed with direct measures, such an observation-induced hunger could be a counterpart to the satiety reported here (appetite shifted by watching alone) and would extend sensory-specific satiety to an observational, social setting, of possible relevance to social eating and to how food imagery affects appetite. Whether observing others eat can genuinely increase hunger and intake remains to be tested with graded hunger measures, a non-eating social control, and objective intake.

### 4.3. Cross-species translation of the animal model

The study parallels, in humans, one aspect of this animal model: in rodents, the value of an ordinary diet shifts with the preference of the foods experienced around it, a cognitive, experience-dependent modulation of food value (Cheng et al., 2024). Watching food being eaten suppressed wanting for the eaten foods, with greater satiety when seen as one’s own, whereas watching another induced less satiety. Building on that principle, the present study shows in humans that food value can be updated through watching eating alone, without any intake. This depended on perspective: self-referential viewing recruited reward- and value-related representations, whereas viewing another eat additionally engaged social-cognitive systems. The human findings thus carry the animal model’s central insight, that food value is experience-dependent, into a social and observational setting. These are now specified at the level of human brain systems: striatal and medial-prefrontal value circuitry for one’s own eating, versus a temporoparietal–prefrontal mentalizing network for another’s. This convergence indicates the video task captures a genuine satiety process rather than a task-specific artifact.

### 4.4. Limitations and future directions

Several limitations qualify these conclusions. The clearest neural effects, the right temporoparietal junction and the striatal value update, survived whole-brain correction, whereas the self-specific valuation effect was supported only at the a-priori level and the satiety-rating signals were weak, so the study was likely under-powered for these. Because the self and other conditions both showed eating and differed only in viewpoint, the design cannot fully separate social attribution from food-specific processing on the other-eating side, and a non-social, non-food control, as a reviewer helpfully suggested, is the more decisive next test. Vertical inversion and the instruction to imagine the eating as one’s own also confound perspective with low-level and agency cues, although converging evidence favors a mentalizing over an imagery or attention account. Appetite was measured from ratings, and a separate experiment with real eating reproduced the key food-specific effect but without dose control. All foods, moreover, were palatable, familiar and highly wanted, which limits generalization and leaves food value partly entangled with familiarity, and the brain-behavior and appetite-state correlations are exploratory. We regard these as boundaries on interpretation rather than threats to the core findings, which rest on convergent behavioral and neural evidence; the individual caveats and the analyses that bear on them are detailed in the Supplement (Extended limitations).

## 5. Conclusions

Watching a short video of food being eaten can change appetite without any eating, and the direction depends on perspective. Watching food being eaten selectively reduced wanting for the eaten foods while leaving non-shown foods more wanted. When the eating was seen as one’s own, this was accompanied by greater satiety and by engagement of food-value and reward circuitry (ventral striatum and medial prefrontal cortex). Watching another eat the same food produced less satiety. It instead engaged the social-cognitive (mentalizing) network centered on the temporoparietal junction, and whether it can itself heighten appetite remains open. Observational sensory satiety thus reflects an update of the motivational value of one’s own eating, dissociable both behaviorally and neurally from the social evaluation of another’s eating. Watching eating, and whose eating it is, can thus change appetite through separate food-value and social-cognitive routes. Because the manipulation is a simple inversion of one video, the paradigm offers a controlled, inexpensive and scalable tool for studying, and potentially influencing, human appetite and food-preference change.

## Ethical statement

The study was approved by the General Ethics Committee of Fukushima Medical University (approval no. REC2023-116) and was conducted in accordance with the Declaration of Helsinki. Participants in the fMRI study (Study 2) gave written informed consent before taking part. The online survey (Study 1) was administered through a research panel operated in accordance with the ESOMAR international code for market, opinion, and social research; panel members provided informed consent electronically before participating, and the investigators analyzed only de-identified responses.

## CRediT authorship contribution statement

**Juri Fujiwara:** Conceptualization, Investigation, Formal analysis, Writing – original draft, Writing – review & editing. **Hitoshi Kubo:** Investigation, Writing – review & editing. **Satoshi Kida:** Conceptualization, Funding acquisition, Supervision, Writing – original draft, Writing – review & editing.

## Data availability

Unthresholded whole-brain statistical maps are publicly available on NeuroVault (https://neurovault.org/collections/24249/). De-identified behavioral data for all studies are publicly available on the Open Science Framework (https://osf.io/wa659/). The video and photograph stimuli are available from the corresponding author upon reasonable request, subject to the consent given by the individual who appears in the videos.

## Acknowledgements

We thank Satoshi Takeda, who appeared as the hand model in the video stimuli and assisted with the real-eating experiment carried out as part of his undergraduate research.

## Supplementary Methods and Results: real-eating experiment

To test whether the wanting-specific satiety observed with the videos also follows actual consumption, six participants (one of whom also took part in the imaging study) completed a real-eating version of the rating task. Inside the scanner they rated 12 snack foods for liking and for wanting (6-point scales). They then left the scanner and, for five minutes, ate as much as they wished of two foods drawn from one taste group (the eaten foods); the remaining two foods of that group served as similar-tasting controls, and the foods of the other groups as different-tasting controls. Each food was provided in its standard commercial retail package; portions were not standardized by weight or energy, and the amount available exceeded what could be consumed in five minutes, so intake reflected ad libitum eating within a fixed window rather than a fixed or exhausted dose. They then re-entered the scanner and re-rated all foods. Change scores (post − pre) were compared across conditions with paired-samples *t*-tests with Holm–Šidák correction (n = 6). Wanting decreased selectively for the eaten foods (Δ = −1.67 ± 0.85) and, to a lesser degree, for similar-tasting foods (Δ = −0.62 ± 0.73), whereas different-tasting foods were unchanged (Δ = +0.17 ± 0.21). This graded ordering reproduces the pattern seen in the main study; with only six participants, however, the individual pairwise differences did not survive correction (eaten vs similar *t*(5) = −2.64, uncorrected p = .046, corrected p = .13; eaten vs different *t*(5) = −2.30, uncorrected p = .069). Liking did not differ across conditions (all p > .3). Actual eating thus reproduced the same wanting-specific, taste-graded pattern seen after watching oneself eat (Figure S6); given the small sample, these findings are reported as convergent, pattern-level support rather than as a powered test.

## Model-comparison and connectivity controls

Two control analyses confirmed the specificity of the main findings. First, we checked whether the results depended on how the eating videos were modeled. We compared our simple model (model 4, which just labels each video as self- or other-eating) with more complex versions that also let each trial’s response scale with how much the food was valued or how full the person felt. The simple model won: adding value scaling fit worse in every one of the 41 participants (BIC −88.4 versus −81.3, where lower is better), and adding satiety scaling made no difference (*p* = 0.46), justifying the event-based models used throughout. Second, we asked whether the mentalizing region (right TPJ) actively communicates with the value regions during the task, using a connectivity analysis (a psychophysiological interaction, model 6). It did not. There was no such task-related coupling either while watching the videos or while rating fullness. The conclusions therefore rest on the consistent activation results, not on directed connectivity between regions.

## Extended limitations

The effects differed in the level of correction they reached. The right temporoparietal junction and the post-consumption striatal and parietal value update survived whole-brain cluster-level correction, whereas the self-specific valuation effect was established at the a-priori small-volume level, and no gustatory or bodily signal reached significance during the satiety rating. The study may have been under-powered for these weaker signals, and larger samples with permutation-based inference would help.

Appetite was assessed from ratings rather than intake. A separate experiment in which participants actually ate the food reproduced the same wanting-specific, taste-graded reduction, linking the effect to real consumption. There, foods were offered in their standard retail package rather than in weight- or energy-matched portions, and participants ate ad libitum for five minutes without exhausting the available amount, which precludes dose-response inference. Even so, the decrease in wanting was steepest for the eaten food, intermediate for similar-tasting foods, and absent for different-tasting foods, tying it to the sensory properties of the eaten food rather than to general gastric fullness. Weight- or calorie-controlled portions would allow a more quantitative test, and we did not measure subsequent intake, physiological markers, or longer-term preference change, so the ecological consequences remain open.

Vertical inversion alters low-level visual and biological-motion cues and perceived agency, so part of the self versus other difference could in principle reflect these rather than perspective, and the perspective cue was also an instruction to imagine the eating as one’s own or another’s. Three observations nonetheless indicate that participants represented observed eating rather than generating unsupported imagery: the footage was real, participants reported perceiving the inverted videos as another person eating, and the other > self contrast engaged the mentalizing network rather than only visual areas, which inversion alone would not predict. Instructed-perspective or matched inverted non-eating motion controls would isolate perspective more fully.

The neural contrast set self against other with both conditions depicting eating, so it cannot separate social attribution from food-specific processing on the other-eating side, and no non-social, non-food baseline was included. A control that involves neither eating nor a social agent, such as watching an object being moved or taken away, would confirm whether food-value regions genuinely track another’s eating; such a control would likely share the domain-general social component but would not be expected to reproduce the food-specific change in appetite, which was selective to the foods shown being eaten and, with real eating, graded by taste. The self > other contrast may likewise reflect in part domain-general self-attribution, since the design lacks a non-eating self/other control; the behavioral outcome, however, was food- and sensory-specific, as self-attribution selectively devalued the eaten foods and updated their value in the ventral striatum, which a generic self-attribution signal could not produce. The eating hand belonged to a single male model, so first-person identification may have been weaker for female participants, although the self condition was defined by viewpoint and instruction rather than by whose hand appeared.

Fasting status was self-reported against a 2-h criterion rather than objectively verified. Because every stimulus depicted food, the design could not isolate food-specific from food-general signals; this is the most likely reason the food-broad insula did not emerge in any differential contrast, and the temporal rise in orbitofrontal and striatal signal during the video, though consistent with accumulating food value, could also reflect sustained attention or anticipation, so a non-food video would be needed to attribute it specifically to food value. The foods were also highly familiar and each participant’s currently most-wanted, so food value is entangled with familiarity, recognition and prior positive experience; varying familiarity, prior experience and novelty would isolate food value from memory-related processing and provide a closer human parallel to the animal model (Cheng et al., 2024). The palatable Japanese snack set may limit generalization across foods and cultures, although the sensory-specific decrease in wanting did not differ by taste or, with taste fixed, by color (Study 1). Finally, the fMRI sample was young, but the survey (ages 15 to 97) showed the behavioral effects generalize across a broad age range, and the brain-behavior and appetite-state correlations are exploratory and await independent replication.

## Supplementary figure legends

**Figure S1.**
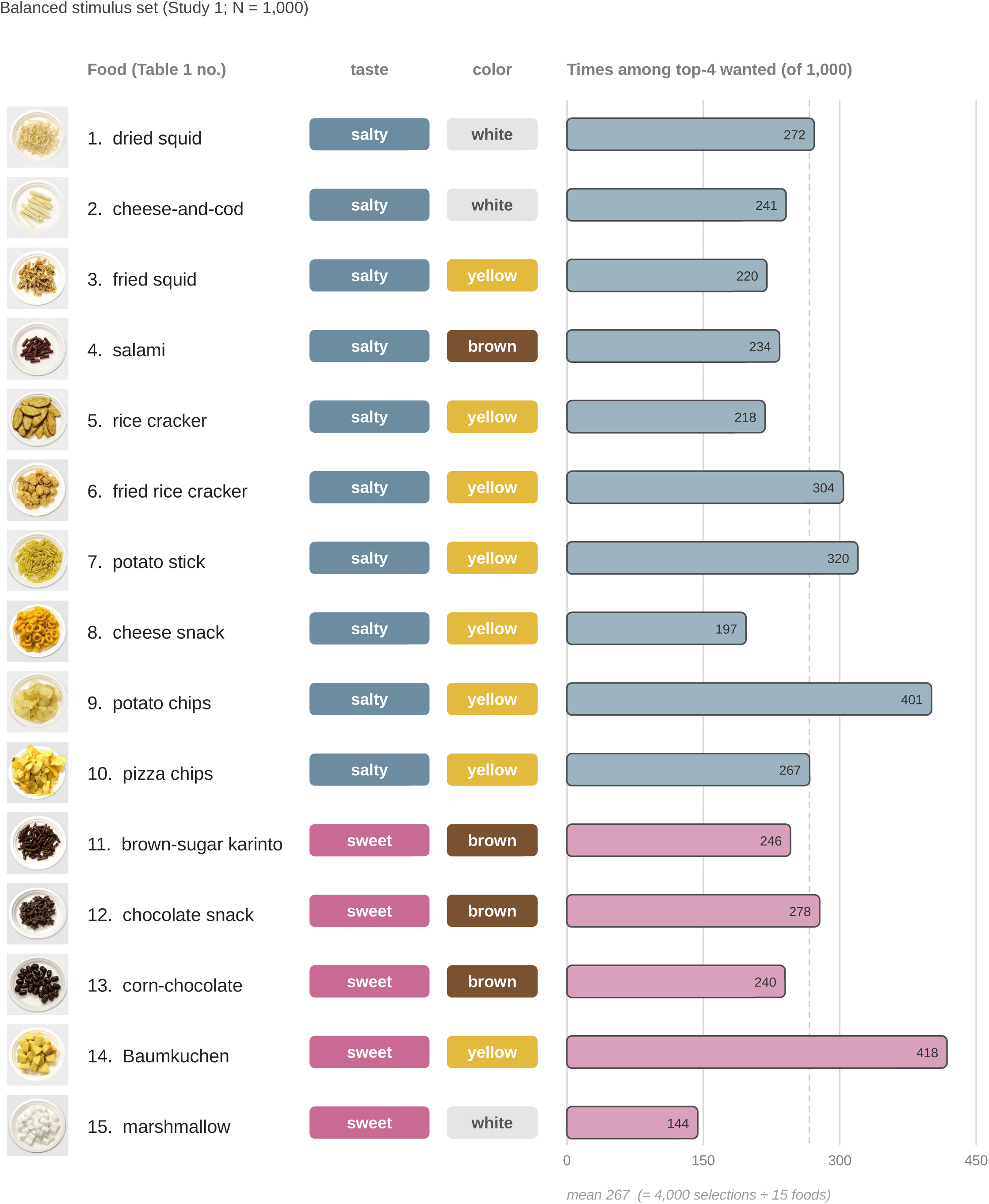
Balanced stimulus set (Study 1; N = 1,000). The set comprised everyday snack foods widely available in Japan, assembled to span the salty and sweet ranges and a range of appearances so that every participant could find foods they genuinely wanted. The 15 snack foods, numbered as in Table 1 (1, dried squid; 2, cheese-and-cod; 3, fried squid; 4, salami; 5, rice cracker; 6, fried rice cracker; 7, potato stick; 8, cheese snack; 9, potato chips; 10, pizza chips; 11, brown-sugar karinto; 12, chocolate snack; 13, corn-chocolate; 14, Baumkuchen; 15, marshmallow), each shown with its taste (salty for 1–10, sweet for 11–15) and a descriptive color label (grouped as yellow/white versus brown for the color-robustness analysis in Figure S2e,f), together with how often it was among the four foods a participant rated highest for wanting. Each food was chosen by 144–418 of 1,000 participants (dashed line, mean); the counts sum to 4,000 (= 1,000 × 4 per participant), and their narrow range indicates a well-distributed, non-idiosyncratic stimulus set.

**Figure S2.**
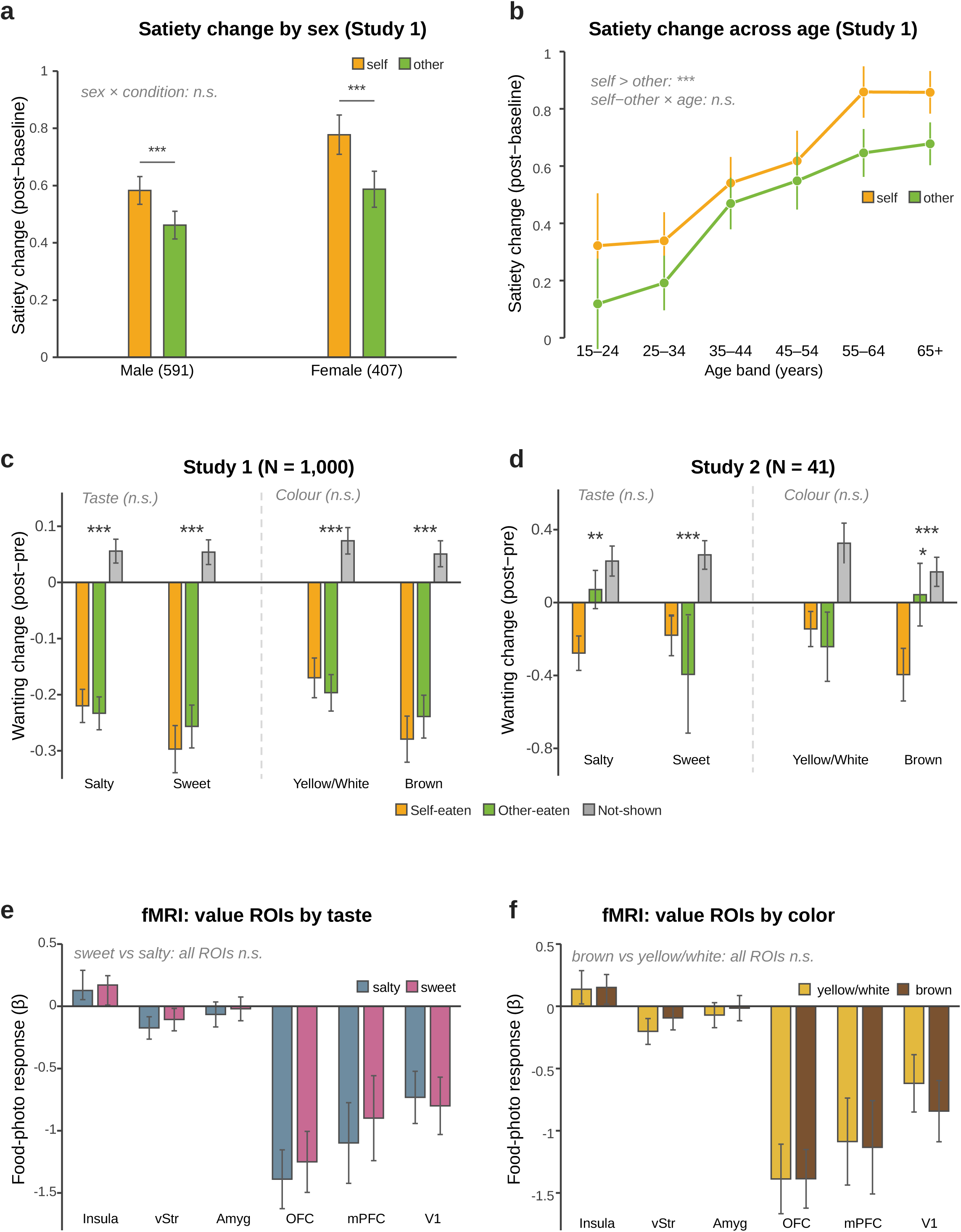
Robustness of the behavioral effects across sex, age and food category. (a) Satiety change (post-video − baseline) for self versus other by sex (Study 1, N = 1,000). (b) Satiety change across age bands (Study 1), for self and other. (c) Study 1: wanting change (post − pre) for self-eaten, other-eaten and not-shown foods, shown separately by taste (salty/savory vs sweet) and by color (yellow/white vs brown). (d) The same breakdown in the fMRI sample (Study 2; N = 41). (e, f) Mean activation to food photos during the pre-run in a-priori food-value ROIs (insula, ventral striatum [vStr], amygdala [Amyg], OFC, mPFC) and a visual control (V1), for (e) salty vs sweet and (f) yellow/white vs brown foods. In (c) and (d), stars above each category test the eaten-versus-not-shown wanting change (Holm–Šidák-corrected). Self, orange; other, green; not-shown, gray. Bars/points, mean; whiskers, s.e.m. \*\*\**p*<.001, \*\**p*<.01, \**p*<.05; n.s., not significant.

**Figure S3.**
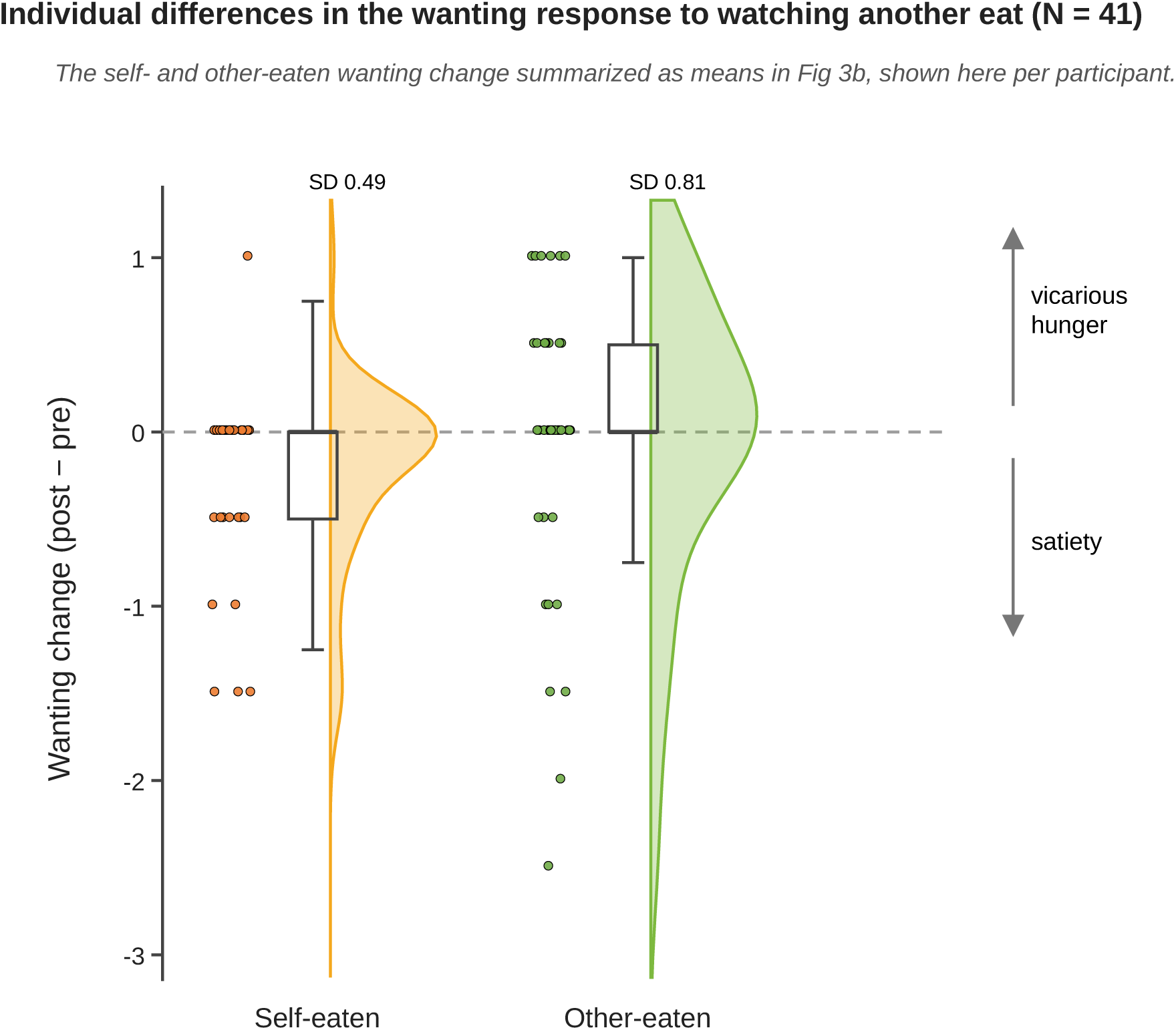
Individual differences in the wanting response to watching another eat (N = 41). Wanting change (post − pre) for the self- and other-eaten foods, shown as a raincloud: one point per participant, with a half-violin density estimate, a box (median and interquartile range) and whiskers.

**Figure S4.**
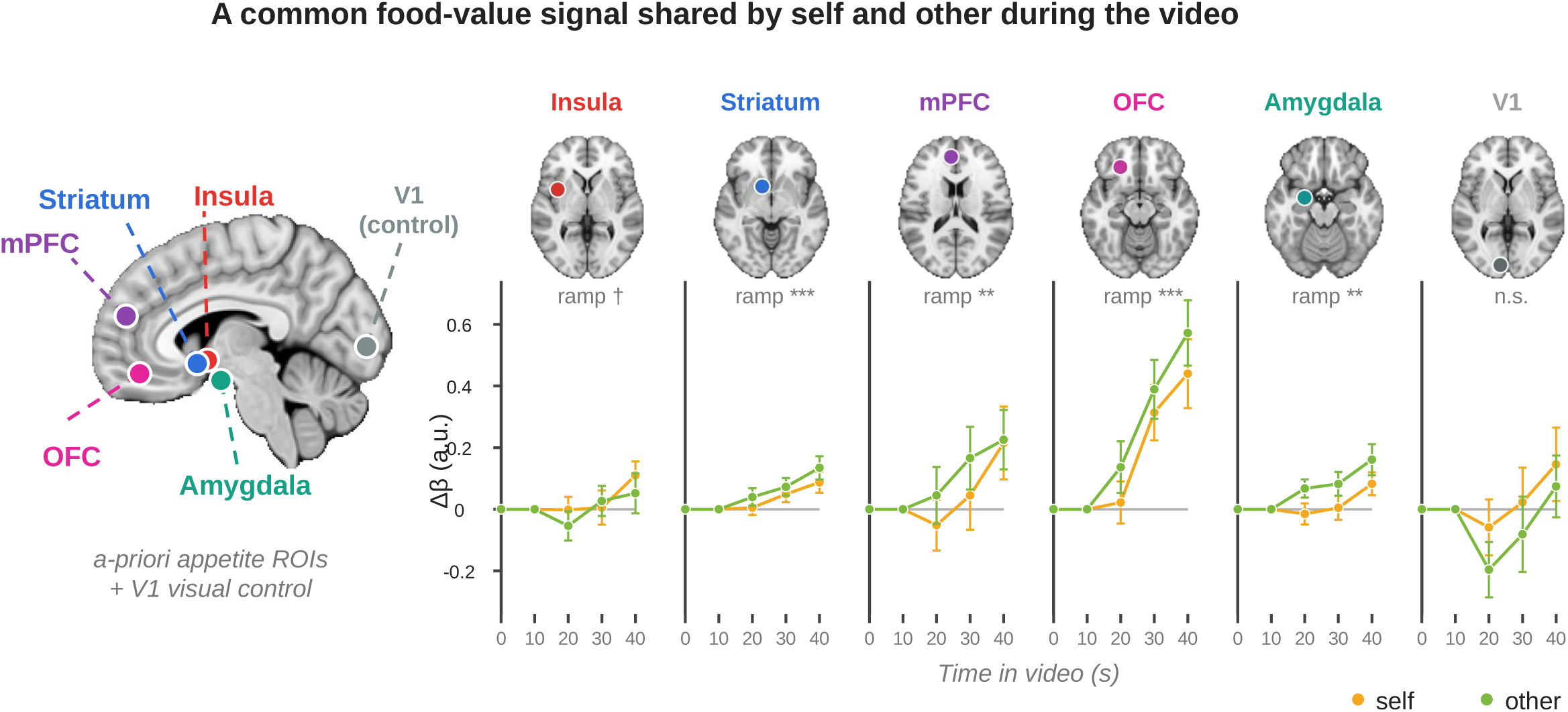
A food-value signal common to self and other during the video (N = 41). Left, the five a-priori appetite ROIs (insula, ventral striatum, amygdala, orbitofrontal and medial prefrontal cortex) together with a V1 visual control, shown as 8-mm spheres on a left-hemisphere sagittal section (MNI152 template, anterior-left convention). Right, BOLD time-courses (Δβ, change from video onset) across the 40-s video (x-axis, seconds from onset) for each region. Activity increased across the video in every appetite region (linear ramp *p* < .01 in all except the insula, *p* = .056) and did not differ between watching oneself and watching another (self versus other n.s.), indicating a common food/eating signal irrespective of whether it was seen as one’s own or another’s; the V1 control showed no such ramp. Self, orange; other, green; lines and points, mean ± s.e.m.

**Figure S5.**
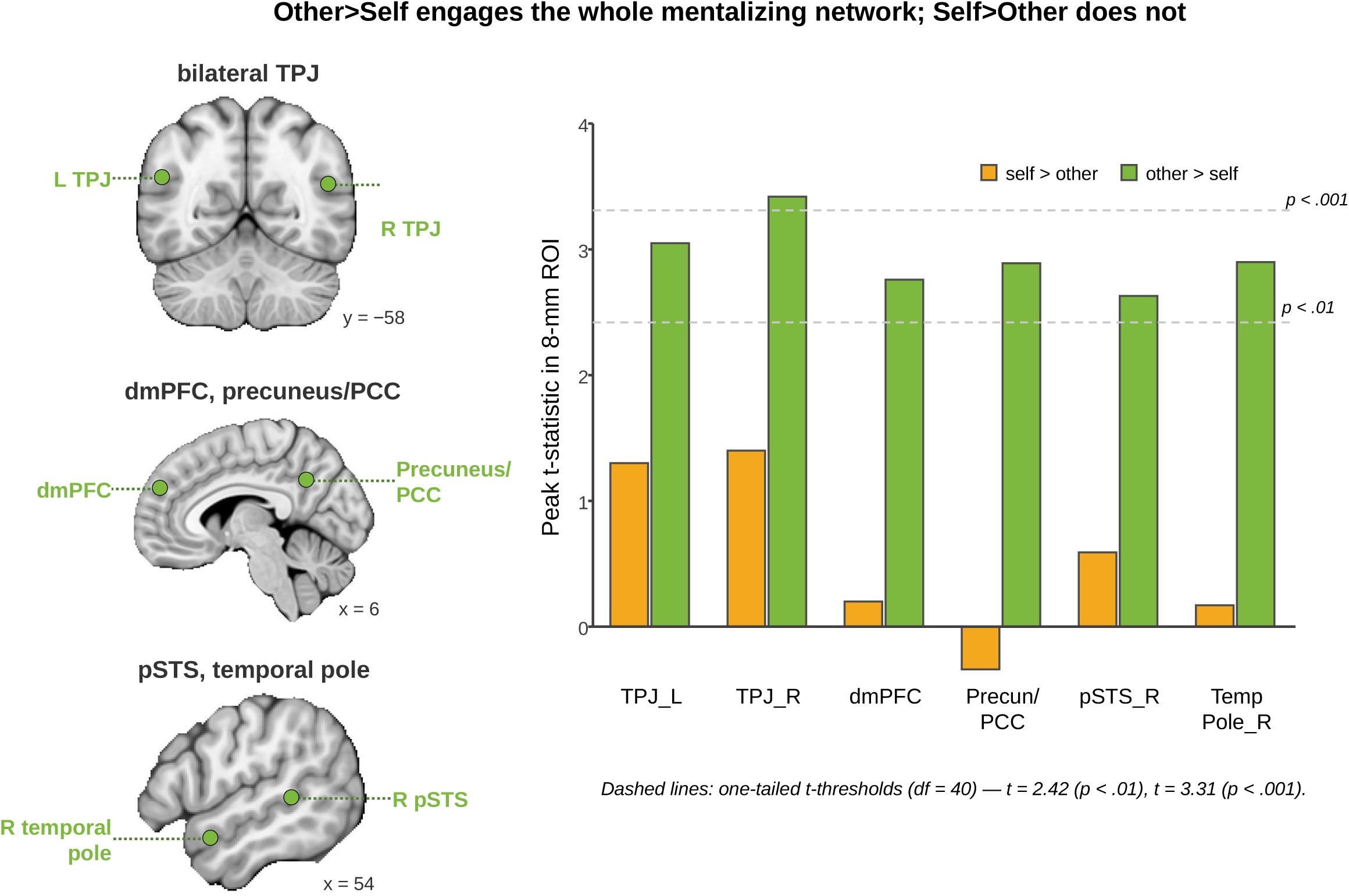
Mentalizing network engaged by watching another eat (N = 41). Peak t within each 8-mm ROI for the other > self and self > other contrasts. Watching another eat (other > self) engaged the whole mentalizing network (bilateral temporoparietal junction, dorsomedial prefrontal cortex, precuneus/posterior cingulate, right posterior superior temporal sulcus and right temporal pole), whereas self > other was null throughout. The brain insets show the ROI locations: a coronal section for the bilateral TPJ, a midline sagittal section for dmPFC and precuneus/PCC, and a lateral sagittal section for the right pSTS and temporal pole (ROIs are 8-mm spheres). Other > self, green; self > other, orange. Dashed reference lines mark one-tailed t-thresholds (df = 40): *t* = 2.42 (p < .01) and *t* = 3.31 (p < .001).

**Figure S6.**
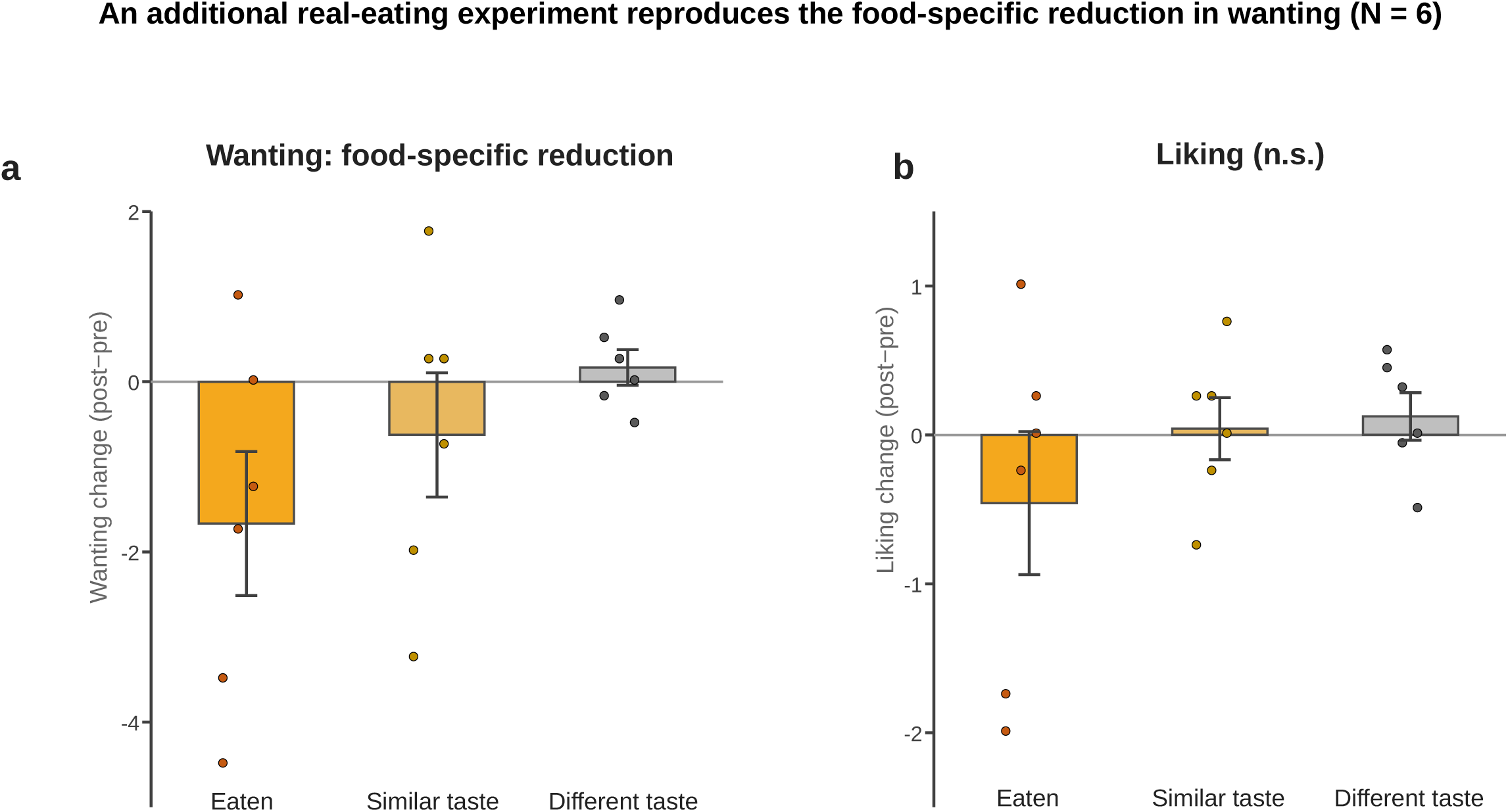
An additional real-eating experiment reproduces the sensory-specific decrease in wanting (n = 6). Six participants (one of whom also took part in the imaging study) rated 12 snack foods for liking and wanting, then left the scanner and actually ate, for five minutes, two foods drawn from one taste group (the eaten foods); afterwards they re-rated all foods. (a) Wanting change (post − pre) decreased selectively for the eaten foods and, to a lesser degree, for similar-tasting foods, whereas different-tasting foods were unchanged. The graded ordering reproduces the main study; with only six participants, the pairwise differences did not survive Holm–Šidák correction (eaten vs similar uncorrected *p* = .046, corrected *p* = .13). (b) Liking change did not differ across conditions. Bars, mean ± s.e.m.; points, individual participants.

